# The Moa the Merrier: Resolving When the Dinornithiformes Went Extinct

**DOI:** 10.1101/2023.08.07.552261

**Authors:** Floe Foxon

## Abstract

The Moa (Aves: Dinornithiformes) are an extinct group of the ratite clade from New Zealand. The overkill hypothesis asserts that the first New Zealand settlers hunted the Moa to extinction by 1450 CE, whereas the staggered survival hypothesis allows for Moa survival until after Europeans began to arrive on New Zealand. Alleged Moa sightings post-1450 CE may shed light on these competing hypotheses. A dataset of 97 alleged Moa sightings from circa 1675 CE to 1993 CE was constructed, with sightings given subjective quality ratings corresponding to various statistical probabilities. Cumulative probabilities of Moa persistence were calculated with a conservative survival model using these probabilistic sighting-records; a method recently applied to sightings of the Thylacine. Cumulative persistence probability fell sharply after 1408 CE, and across pessimistic and optimistic variations of the model, it was more likely than not that the Moa were extinct by 1770 CE. Probabilistic sighting-record models favour the overkill hypothesis, and give very low probabilities of Moa persistence around the time of European arrival. Eyewitness data on Moa sightings are amenable to scientific study, and these methods may be applied to similar animals.

## Introduction

In a recent study in *Science of the Total Environment*, Brook, Sleightholme, et al. (2023) used a database of alleged Thylacine sighting records from 1910 CE to 2023 CE in a computational extinction dynamics estimator (EDE) to estimate probabilities of persistence of the predatory marsupial Thylacine, *Thylacinus cynocephalus*. Brook, Sleightholme, et al. (2023) bring a much-needed level of nuance, scientific rigour, and statistical methodology to bear on a topic that is often shrouded in pseudoscience and incredulity; by assigning probabilities to individual sightings rather treating these in binary, Brook et al. transcend the traditional paradigm of either blind belief in, or arrogant denial of, the existence of hidden animals.

The methods presented by Brook, Sleightholme, et al. (2023) may be applied to other species whose extinction dates are contested. Roughly 2,000 kilometers east of the Thylacine’s native Tasmania lies New Zealand, a land dominated by avifauna, the greatest among which were the Dinornithiformes or Moa: large flightless birds of the ratite clade (Worthy and Holdaway, 2002). Moa lived in relative peace on New Zealand as megaherbivorous browsers (Wood et al., 2020), and were preyed upon only by Haast’s eagle, *Harpagornis moorei* (Brathwaite, 1992; Worthy and Holdaway, 2002), until the arrival of the Austronesian peoples circa 1250–1300 CE (Irwin and Walrond, 2016).

The latest and now generally-accepted view is that, as part of the greater Quaternary ‘overkill’ (Sandom et al., 2014), an extremely low-density population of first New Zealand settlers wiped out the entire population of Moa on the islands in a period of less than 150 years (Holdaway, Allentoft, et al., 2014; Holdaway and Jacomb, 2000), which would coincide with 1400–1450 CE. This rapid megafaunal extinction is evidenced by high-quality radiocarbon dates on Moa remains from natural and archaeological sites (Allentoft et al., 2014; Holdaway, Allentoft, et al., 2014; Perry et al., 2014). Indeed, it is thought that the Moa had relatively low reproduction rates and older ages of sexual maturity (Turvey, Green, et al., 2005; Turvey and Holdaway, 2005) making them vulnerable to extinction via predation, and a recent collaborative analysis of ancestral sayings suggests that early Maori used the Moa as a metaphor for extinction, e.g. ‘lost as the Moa is lost’, similar to the English phrase ‘dead as a dodo’ (Wehi et al., 2018).

The overkill hypothesis was not a view held by all; in particular, critics pointed to the marked variability in terrain across New Zealand, and questioned the ability of settlers to eradicate all Moa species across a vast, difficult, and (then) unknown land at such speed (Diamond, 2000). Rather than a rapid war against the Moa, critics opted for ‘staggered survival’, perhaps until circa 1800 CE (Silverberg, 1973, Chapter 7), by which time Europeans had begun to arrive on the islands. Some (previous) critics are now in support of the overkill hypothesis (Holdaway, 2023).

Entertaining the possibility of Moa persistence post-1450 CE, which the model by Holdaway, Allentoft, et al. (2014) does not completely preclude, when did the Moa go extinct? Like the Thylacine studied by Brook, Sleightholme, et al. (2023), alleged Moa sightings have persisted to as recently as 1993 CE (Spittle, 2010), with reports originating from both Maori and European peoples (Silverberg, 1973, Chapter 7). Other works have discussed this possibility, including Heuvelmans (1986) which lists “A surviving species of the Moa family… known to some as *roa-roa*” in an ‘Annotated Checklist of Apparently Unknown Animals,’ as well as Mackal’s (1983) work on ‘hidden animals’.

The aim of the present study was to extend the methods of Brook, Sleightholme, et al. (2023) to the Moa as a test of the ‘staggered survival’ hypothesis, i.e. that the Moa persisted long after 1450 CE, by which time the competing ‘overkill’ hypothesis asserts that the Moa were extinct.

## Material and methods

### Data

A new dataset named the New-Zealand Moa Sighting Records Database (NMSRD) was constructed following the Tasmanian Thylacine Sighting Records Database (TTSRD) of Brook, Sleightholme, et al. (2023). Alleged sightings of Moa in New Zealand were taken manually from Spittle (2010) and were coded in a flat-file format (.xlsx) with one observation per row. The columns in this dataset correspond to unique sighting ID; sighting location; year of observation; month of observation; location name; subjective quality-rating (a score between 1 [lowest] to 5 [highest]); number of human observers; number of Moa; reference; observer name(s); and sighting description.

Following Brook, Sleightholme, et al. (2023), sightings rated 5 were assigned a probability of 1; 4-rated sightings were assigned a probability of 0.05; 3-rated sightings 0.01; 2-rated sightings 0.005; and 1-rated sightings 0.001.

For clarity and transparency on the subjective rating system, the following rules were generally applied.

1. Sightings that were quite possibly hoaxes, sightings that were almost certainly cases of misidentification, sightings with reference to the informant’s ‘father’ or ‘grandfather’ (which may be a mistranslation of ‘ancestors’), sightings that referred to ‘fresh’ looking bones (which may have been well-preserved and therefore deceptively old), and especially those with uncertainty in the number and identities of the observers, the date and location of the observation, &c, were generally rated 1 (the lowest rating).
2. Sightings that were less likely to be hoaxes and less clearly misidentification, especially those with known observer identity/date/location/&c, were generally rated 2.
3. Sightings with circumstantial evidence in support (e.g., bones subsequently found in a location where moa were reportedly hunted recently, reported moa behaviour consistent with other struthious birds or reported appearance consistent with museum specimens unknown to the observer, and tracks of uncertain age observed by an expert) were generally rated 3.
4. Sightings involving direct observation by an expert zoologist or ornithologist were rated 4.
5. Radiocarbon dated Moa specimens described in Supplementary Table 1 of Holdaway, Allentoft, et al. (2014) from archaeological sites across New Zealand (i.e., Moa remains contemporaneous with people in New Zealand) were rated 5. These were included because the model requires starting values for which Moa persistence was definite.

Confirmed hoaxes were necessarily excluded.

## Model

With each individual sighting applied a rating (and corresponding probability) as described above, the approach taken to model these sightings was a frequentist approach involving only probabilistic sighting-records and no EDE. This was the most conservative (i.e., pessimistic) survival model applied by Brook, Sleightholme, et al. (2023), resulting in the lowest persistence probabilities in that study. The model is described in a previous publication (Brook, Buettel, et al., 2019), and is summarised succintly as follows:

1. The probability that there was at least one true sighting (i.e. Moa persistence) in year *j*, or the ‘combined’ probability for year *j*, is given by

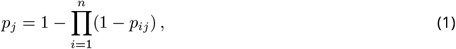

where *p*_*ij*_ is the *i*^th^ sighting in year *j*, and *n* is the total number of sightings in year *j*. For a year with *n* = 1 sighting, this equation intuitively reduces to *p*_*ij*_ = *p*_1_.
2. The cumulative combinatorial probability (denoted by the superscript *c*) of Moa persistence in year *j* is then given by

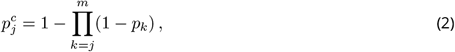

where *p*_*k*_ is the ‘combined’ probability for year *k* (from step 1. above) and *m* is the total number of years for which there are sightings.
3. For variance estimation, jackknife resampling was applied, which effectively tests the sensitivity of the model to any one sighting. This involved removing (with replacement) each sighting in turn and rerunning the model to obtain replicate values for *p*^*c*^ in each year, denoted *p*^*c*^ . The jackknife mean probability for each year was estimated as

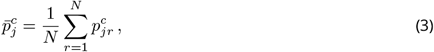

where *N* is the total number of replicates (i.e. the total number of sightings across all years). The variance for each year is estimated as

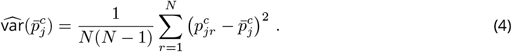

A conservative modelling approach is justified here because the sightings in the NMSRD were generally of lower quality than those in the TTSRD (e.g., including accounts of second or greater hand) and because the present study extends the method in time (circa 1675 CE to 1993 CE here, vs 1910 CE to 2019 CE in Brook, Sleightholme, et al. (2023)), which is disputable (Holdaway, 2023).

### Conservative Measures and Sensitivity Tests

In addition to the most pessimistic survival model in Brook, Sleightholme, et al. (2023) being used, further conservative measures taken were as follows.

- Related sightings (e.g., multiple sightings made by the same individual) were aggregated into single records in the dataset, which had the effect of decreasing the cumulative probability in a given year.
- To test the sensitivity of the model to the subjective probability assigned to each sighting, a ‘pessimistic’ model assigning a probability of 0.001 to all sightings rated 1 to 4, an ‘optimistic’ model assigning a probability of 0.005 to all sightings rated 1 and 2, and a ‘super optimistic’ model assigning a probability of 0.01 to all sightings rated 1 to 3, were implemented (all other probabilties kept the same as in the main model).
- As additional sensitivity tests, sightings of Moa surface remains (i.e. bones, skin, and feathers which may have been deceptively old and well preserved), and sightings of second or great hand were removed from the dataset separately to assess their impact on the results.
- Where a range of years was given for a sighting (e.g., 1800–1830 CE) the approximate midpoint was taken (e.g., 1815 CE), except where a preferred year was provided in the source text.
- Chapters/sightings in Spittle (2010) in which the persistence of Moa into recent times was not explicitly asserted or implied by the observer, as well as sightings for which no specific date or range of dates were provided or could be reasonably deduced (e.g. in the “dim past” or “olden times”), were all excluded from the dataset.

Analyses were perfomed in Python version 3.8.16 with the packages Numpy 1.21.5, Pandas 1.5.2, and Matplotlib 3.6.2. The NMSRD and analysis code are made freely available in the online supplemental materials at https://doi.org/10.17605/OSF.IO/E59H6.

## Results

The NMSRD consists of 96 radiocarbon-dated Moa specimens and 97 alleged sighting records involving a total of over 160 eyewitnesses. The NMSRD contains 66 sightings rated 1, 26 sightings rated 2, five sightings rated 3, zero sightings rated 4, and 96 radiocarbon dated Moa specimens rated 5. Sighting dates ranged from circa 1675 CE to 1993 CE.

The cumulative combinatorial probability of persistence of the Moa over time is plotted in Figure 1, where a value of 1.0 implies the Moa were definitely extant at that time, and a value of 0.0 implies the Moa were definitely extinct at that time. Cumulative combinatorial persistence probability fell sharply after 1408 CE (the last radiocarbon dated Moa specimen in the dataset). For 1770 CE, the cumulative persistence probability was 0.21 in the main model, 0.09 in the pessimistic model, 0.39 in the optimistic model, and 0.61 in the super optimistic model. Therefore, in all but the most optimistic model, the cumulative persistence probability was *<* 0.5 circa 1770 CE. This may be interpreted as it being more likely than not that the Moa were extinct circa 1770 CE. Results are practically unchanged by removing the earliest three records from 1675 CE to 1725 CE. All models provided cumulative persistence probabilities *≤* 0.17 circa 1900 CE and *≤* 0.01 circa 2000 CE.

**Figure 1.**
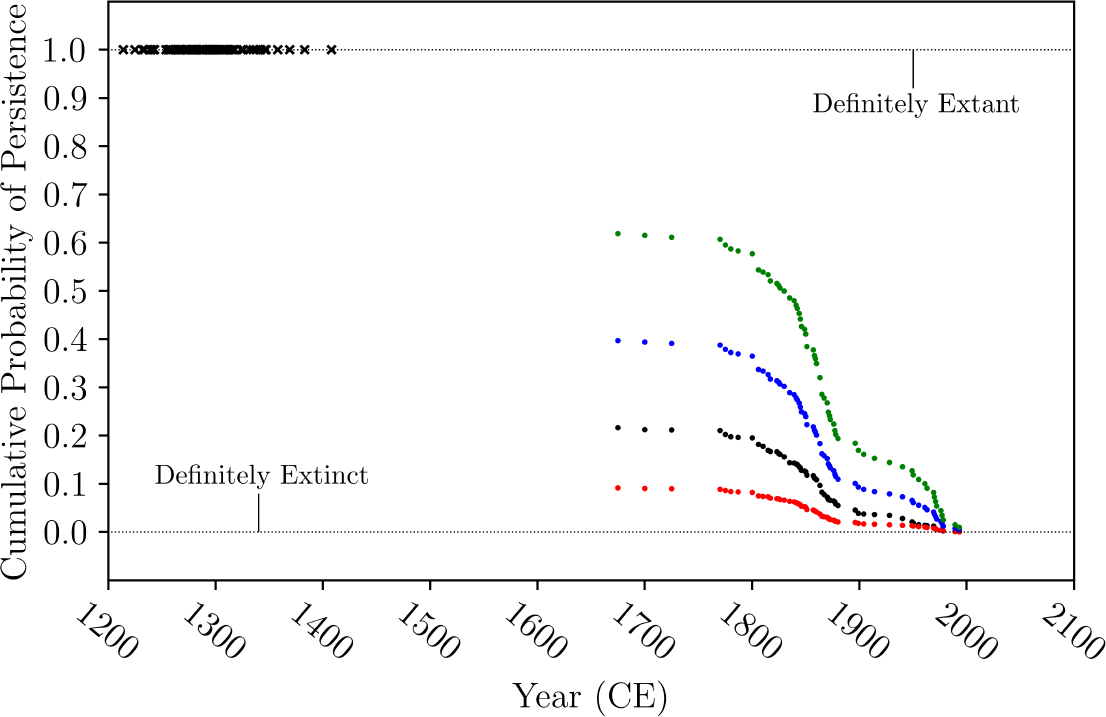
Cumulative probability of Moa persistence over time, as inferred from alleged eyewitness Moa sightings in New Zealand. Dots represent Moa sightings from Spittle (2010), combined in each year. Red dots represent the ‘pessimistic’ model, black dots the main model, blue dots the ‘optimistic’ model, and green dots the ‘super optimistic’ model (see Methods). Black crosses represent radiocarbon dated Moa specimens from archaeological sites (Holdaway, Allentoft, et al., 2014). Results are practically unchanged by removing the earliest ‘sightings’ circa 1675 CE to 1725 CE.

On model sensitivity, jackknife resampling estimates for the cumulative combinatorial persistence probabilities differed from non-resampling results only by a few thousandths (e.g. 0.213 vs 0.211), and variance estimates derived from jackknife resampling were at most of the order 10^*−*6^. These resampling results suggest that the model is not particularly sensitive to any one sighting. Additionally, 20% of reports were of surface Moa remains (bones, skin, and feathers) and more than two thirds were of second or greater hand. Surface remains reports had only a small impact on results, e.g. circa 1770 CE the cumulative persistence probability was 0.21 in the main model with suface remains reports included, and 0.19 with these reports excluded. Secondhand reports, comprising a far greater number of sightings, had a proportionally greater impact on the model. Results of the main model excluding secondhand reports were similar to results of the pessimistic model, e.g. circa 1800 CE the cumulative persistence probability was 0.08 in the pessimistic model with secondhand reports included, and 0.09 in the main model with secondhand reports excluded.

## Discussion

Conservatively assuming a low but non-zero probability of Moa persistence associated with each alleged Moa sighting post-1450 CE, it is more likely than not that the Moa were extinct before 1770 CE, when Europeans began arriving on New Zealand. This finding favours the overkill hypothesis. Moa sightings in the period post-1450 CE are likely not numerous or reliable enough to support Moa survival into more recent times. Only if one assumes the most optimistic model (generously assigning a probability of 0.01 to all of the sightings) does the extinction of the Moa in New Zealand become likely as late as circa 1850 CE. Indeed, there were very few Maori proverbs relating to Moa extinction before 1800 CE (Wehi et al., 2018), which could suggest persistence of some Moa species up to that time.

This work provides for the first time in the published literature an estimate for the extinction date of the Moa on statistical principles using a probabilistic sighting-record model, as in Brook, Sleightholme, et al. (2023). This study is perhaps more nuanced than previous studies, and demonstrates that the methods of Brook, Sleightholme, et al. (2023) are applicable to other controversial animals besides the Thylacine.

The results of the present study are comparable to the Bayesian extinction analyses of Holdaway, Allentoft, et al. (2014). Rather than replacing or surpassing the previous Bayesian model, the sighting record model supports and supplements the previous model with different approach and data. The Bayesian analyses have long tails extending to the present, but with vanishingly small probabilities after 1750 CE. Similarly, the sighting record model has small probabilities circa 1750 CE which vanish toward the present.

Important limitations must be noted. First is the contentious assumption in assigning subjective non-zero probabilities to all moa sightings. Given the lack of recent physical material (i.e., specimens), it is likely that many of the sightings in the NMSRD constitue innocent cases of mistaken identity, or else hoaxes. Nickell (2017) notes that most Moa ‘sightings’ occured around the time when fossil Moa were first described by Richard Owen in 1839 CE (Silverberg, 1973, Chapter 7). This may be evidence of ‘expectant attention’; the tendency for observers to ‘see’ what they anticipate they will see (Radford and Nickell, 2006, Chapter 2). Statistical evidence for expectant attention in the context of other ‘maybe’ animals has been demonstrated by the author in the context of Lake Champlain (Foxon, 2023b). This said, the model was conservative in other ways.

A further limitation is that the NMSRD is not final and is subject to change. The dataset may contain errors that can be ammended as new historical material is brought to the attention of the author. Readers are encouraged to download the data and code in the online supplemental materials and perform their own tests of the subjective ratings/probabilities and adding/removing records as they feel is appropriate. The dataset is also almost certainly incomplete, in part because some ‘sightings’ will go unreported for fear of ridicule. According to William Scoble who reported their alleged Moa sighting to the Dominion Museum director R. A. Falla, Scoble did not want to be met “with the ridicule metered out to those who talk of sea serpents… when one has met a moa [sic] so close at hand and over so many years reads so many conflicting statements from Science one must be forgiven for reminding you how grey can be all theory” (Scoble, 1952).

Another limitation is that this model applies generally to the order Dinornithiformes, and does not distinguish between individual Moa species. Indeed, there were inhomogeneities across sightings in morphological characteristics such as size and plumage (see ‘Description’ column of the NMSRD dataset), which could either suggest different species/sex/ages of the animals being described, or simply noise from misidentification and hoaxes.

Importantly, as noted by Holdaway (2023), the NMSRD contains some highly uncertain sightings, including uncertainties about the age of the purported witness, inconsistencies with known Moa biology (e.g. many eggs or young), and in some cases the profit motive for faking a sighting to enhance business in a given area. For transparency, the author similarly approached these sightings skeptically. Given that removing such sightings would necessarily decrease the persistence probability results, and given that the models presented in this study already favour the overkill hypothesis, it follows that any further filtering of the data would only favour the overkill hypothesis further. Readers may explore these possibilities using the data and code provided.

The reliance in the present study and in Brook, Sleightholme, et al. (2023) on eyewitness testimony (i.e., circumstantial evidence) to infer the existence of an animal is, in a sense, ‘folk zoology’, and represents a more mathematically-grounded implementation of the methods described by Heuvelmans (1982, 1984, 1988) in his seminal works. The Thylacine and Moa are animals of known taxa, believed to have become extinct during historical times, and whose existence in the present is unrecognised in conventional zoology. These are described as ‘Category IV’ animals in the classificatory system of hidden or unknown animals described by ecologist J. Richard Greenwell (1985). Other Category IV animals, such as the ivory-billed woodpecker, *Campephilus principalis*, are amenable to the same probability-based analysis (Brook, Buettel, et al., 2019), and future works may investigate other such animals using these methods.

In conclusion, a probabilistic sighting-record model favours the overkill hypothesis of rapid Moa extinction. Still, eyewitness data on Moa sightings are amenable to meaningful scientific analysis as a form of citizen science, similar to previous studies on sightings of animals in folklore (Brook, Sleightholme, et al., 2023; Foxon, 2023a,b; Paxton, 2009). Regretfully, the Moa are now extinct. As Silverberg (1973, Chapter 7) wrote, “The mighty *Dinornis*… is no more likely to be seen again than Sinbad’s rukh.” The Moa have had a tremendous impact on New Zealand culture (Armstrong, 2010), and doubtless will continue to do so.

## Acknowledgements

The author thanks Drs Werner Ulrich, Richard N. Holdaway, and Tim Coulson for helpful comments on and suggestions for the manuscript. The pun in the title of this article originates from an anonymous comment in the *Otago Witness* column ‘Passing Notes’ from October 10, 1874.

## Data, script, code, and supplementary information availability

Data and code are available online: https://doi.org/10.17605/OSF.IO/E59H6.

## References

Allentoft ME, R Heller, CL Oskam, ED Lorenzen, ML Hale, MTP Gilbert, C Jacomb, RN Holdaway, and M Bunce (2014). Extinct New Zealand megafauna were not in decline before human colonization. Proceedings of the National Academy of Sciences 111, 4922–4927. 10.1073/pnas.1314972111.

Armstrong P (2010). Moa Citings. The Journal of Commonwealth Literature 45, 325–339. 10.1177/0021989410376799.

Brathwaite DH (1992). Notes on the weight, flying ability, habitat, and prey of Haast’s Eagle (Harpagornis moorei). Notornis 39, 239–247.

Brook BW, SR Sleightholme, CR Campbell, I Jarić, and JC Buettel (2023). Resolving when (and where) the Thylacine went extinct. Science of The Total Environment 877, 162878. 10.1016/j.scitotenv.2023.162878.

Brook BW, JC Buettel, and I Jarić (2019). A fast re-sampling method for using reliability ratings of sightings with extinction-date estimators. Ecology 100, e02787. 10.1002/ecy.2787.

Diamond J (2000). Blitzkrieg Against the Moas. Science 287, 2170–2171. 10.1126/science.287.5461.2170.

Foxon F (2023a). If it’s there, could it be a bear? PeerRef. 10.1101/2023.01.14.524058.

Diamond J (2023b). What’s in Lake Champlain? Analysing historic sightings of the cryptid known as “Champ”. The Skeptic. https://www.skeptic.org.uk/2023/06/whats-in-lake-champlain-analysing-historic-sightings-of-the-cryptid-known-as-champ/.

Greenwell JR (1985). A Classificatory System for Cryptozoology. Cryptozoology 4, 1–14.

Heuvelmans B (1982). What is Cryptozoology? Cryptozoology 1, 1–12.

Heuvelmans B (1984). The Birth and Early History of Cryptozoology. Cryptozoology 3, 1–30.

Heuvelmans B (1986). Annotated Checklist of Apparently Unknown Animals With Which Cryptozoology is Concerned. Cryptozoology 5, 1–26.

Heuvelmans B (1988). The Sources and Method of Cryptozoological Research. Cryptozoology 7, 1–21.

Holdaway RN, ME Allentoft, C Jacomb, CL Oskam, NR Beavan, and M Bunce (2014). An extremely low-density human population exterminated New Zealand moa. Nature Communications 5, 5436. 10.1038/ncomms6436.

Holdaway RN and C Jacomb (2000). Rapid extinction of the moas (Aves: Dinornithiformes): model, test, and implications. Science 287, 2250–2254. 10.1126/science.287.5461.2250.

Holdaway RN (Sept. 20, 2023). Personal communication.

Irwin G and C Walrond (2016). When was New Zealand first settled? Te Ara - the Encyclopedia of New Zealand. http://www.TeAra.govt.nz/en/when-was-new-zealand-first-settled.

Mackal RP (1983). Searching for Hidden Animals. Cadogan Books.

Nickell J (2017). The New Zealand Moa: From Extinct Bird to Cryptid. Skeptical Inquirer. https://skepticalinquirer.org/newsletter/the-new-zealand-moa-from-extinct-bird-to-cryptid/.

Paxton CGM (2009). The plural of ‘anecdote’ can be ‘data’: statistical analysis of viewing distances in reports of unidentified large marine animals 1758–2000. Journal of Zoology 279, 381–387. 10.1111/j.1469-7998.2009.00630.x.

Perry GL, AB Wheeler, JR Wood, and JM Wilmshurst (2014). A high-precision chronology for the rapid extinction of New Zealand moa (Aves, Dinornithiformes). Quaternary Science Reviews 105, 126–135. issn: 0277-3791. 10.1016/j.quascirev.2014.09.025.

Radford B and J Nickell (2006). Lake Monster Mysteries: Investigating the World’s Most Elusive Creatures. University Press of Kentucky.

Sandom C, S Faurby, B Sandel, and JC Svenning (2014). Global late Quaternary megafauna extinctions linked to humans, not climate change. Proceedings of the Royal Society B: Biological Sciences 281, 20133254. 10.1098/rspb.2013.3254.

Scoble W (1952). Unpublished letter to Dr R. A. Falla, Director of the Dominion Museum. Dated December 28, 1952.

Silverberg R (1973). The Dodo, the Auk and the Oryx: Vanished and Vanishing Creatures. Puffin Books.

Spittle B (2010). Moa Sightings Volumes 1–3. Paua Press Limited, Dunedin.

Turvey ST, OR Green, and RN Holdaway (2005). Cortical growth marks reveal extended juvenile development in New Zealand moa. Nature 435, 940–943. 10.1038/nature03635.

Turvey ST and RN Holdaway (2005). Postnatal ontogeny, population structure, and extinction of the giant moa Dinornis. Journal of Morphology 265, 70–86. 10.1002/jmor.10341.

Wehi PM, MP Cox, T Roa, and H Whaanga (2018). Human perceptions of megafaunal extinction events revealed by linguistic analysis of indigenous oral traditions. Human Ecology 46, 461–470. 10.1007/s10745-018-0004-0.

Wood JR, SJ Richardson, MS McGlone, and JM Wilmshurst (2020). The diets of moa (Aves: Dinornithiformes). New Zealand Journal of Ecology 44, 3397. 10.20417/nzjecol.44.3.

Worthy TH and RN Holdaway (2002). The Lost World of the Moa: Prehistoric Life of New Zealand. Indiana University Press.

